# Diet-induced obesity impacts influenza disease severity and transmission dynamics in ferrets

**DOI:** 10.1101/2023.09.26.558609

**Authors:** Victoria Meliopoulos, Rebekah Honce, Brandi Livingston, Virginia Hargest, Pamela Freiden, Lauren Lazure, Pamela H. Brigleb, Erik Karlsson, Heather Tillman, E. Kaity Allen, David Boyd, Paul G. Thomas, Stacey Schultz-Cherry

## Abstract

Obesity, and the associated metabolic syndrome, is a risk factor for increased disease severity with a variety of infectious agents, including influenza virus. Yet the mechanisms are only partially understood. As the number of people, particularly children, living with obesity continues to rise, it is critical to understand the role of host status on disease pathogenesis. In these studies, we use a novel diet-induced obese ferret model and new tools to demonstrate that like humans, obesity resulted in significant changes to the lung microenvironment leading to increased clinical disease and viral spread to the lower respiratory tract. The decreased antiviral responses also resulted in obese animals shedding higher infectious virus for longer making them more likely to transmit to contacts. These data suggest the obese ferret model may be crucial to understanding obesity’s impact on influenza disease severity and community transmission, and a key tool for therapeutic and intervention development for this high-risk population.

**Teaser:** A new ferret model and tools to explore obesity’s impact on respiratory virus infection, susceptibility, and community transmission.

## Introduction

Obesity, and the associated metabolic syndrome, is a risk factor for increased disease severity with a variety of infectious agents, including influenza virus, SARS-CoV-2, respiratory syncytial virus (RSV), and bacterial infection (*1–3*). Human cohort studies have reported that participants with a higher body mass index (BMI) are more likely to report symptoms, may have higher viral titers in exhaled breath (*4*, *5*) and prolonged shed time resulting in a longer duration of illness, as well as increased susceptibility to certain strains of influenza virus (*6*, *7*). Yet the mechanisms behind increased disease severity in obese individuals are only partially understood. As the number of people, particularly children, living with obesity continues to rise (*8–11*), it is critical to understand the role of host status on disease pathogenesis.

Genetic and diet-induced obese mouse models (*12–14*) have provided valuable insights into the impact of obesity on influenza disease, viral evolution, and therapeutic strategies (*15–18*). Yet, it is difficult to study many key aspects of disease, including viral transmission, in mice. Ferrets, however, can be infected with circulating human viruses without the need for adaptation, display similar clinical signs as humans, and can transmit virus through direct contact and via aerosols (*19*, *20*). We hypothesized that these animals could provide better clarity as to how obesity could impact influenza dynamics in human populations.

While it is possible to make transgenic ferrets (*21*, *22*), diet-induced obesity is a more accurate representation of human obesity. In these studies, we describe the development of a novel diet-induced obese ferret model and the impact on influenza disease severity and transmission. Obese ferrets have increased waist circumference and additional visceral fat compared to control ferrets fed a standard chow. They also had clinical parameters associated with development of metabolic syndrome (MetS). When infected with influenza virus, obese ferrets experienced more severe clinical symptoms than control ferrets of healthy weight independent of viral strain. Lung injury and viral spread to the lower respiratory tract was enhanced. Further, we designed a new panel of validated primers and probes targeting immune and lung function genes, and found obese ferrets had a dysregulated antiviral response in the lungs at baseline and post-infection.

Diet also influenced viral transmission. Obese ferrets were more likely to transmit an avian-like H9N2 influenza virus to other obese ferrets, while likelihood of transmission was equivalent when the transmitting ferret was on our control diet. Overall, our studies show obesity in ferrets impacts disease pathogenesis and transmission. This model, and the newly developed tools, will be invaluable for better understanding the impact of obesity on disease pathogenesis, viral transmission and evolution. and will be crucial for assessing the impact of vaccines and antivirals on disease severity within this increasing population.

## Results

### Inducing obesity and associated metabolic syndrome in ferrets

Male ferrets were maintained on a high calorie diet (obese) or a nutritionally balanced calorie restricted diet (control) for a period of 12 weeks as described in Table S1. Ferrets in the obese group gained significantly more weight (Fig. S1A). Waist circumference, which is strongly associated with increased risk of mortality in humans (*23*, *24*), abdominal adipose tissue, plasma leptin levels, skinfold fat, and overall body mass index were elevated in the obese group (Fig. S1B-F). At the conclusion of the diet period, obese ferrets had developed biochemical hallmarks of metabolic syndrome (MetS) such as increased fasting glucose and total cholesterol (Fig. S1G-H). High-density lipoprotein (HDL) was reduced in obese ferrets while low-density lipoprotein (LDL) was increased, indicating an unhealthy cholesterol balance (Fig. S1I-K), and are clinically associated with human obesity (*25*).

Obesity has been shown to impact the lung microenvironment and is associated with increased expression of pro-inflammatory cytokines and chemokines (*26*, *27*). Because the availability of ferret reagents is limited, we designed a panel of primers targeting lung function and immune response genes (Table S2) and validated as described in methods. We normalized gene expression in the uninfected obese ferrets to the corresponding gene expression in the uninfected control to compare status of the lung microenvironment at baseline (Fig. 1A-B). Even at baseline, gene expression differed in obese compared to control ferrets; globally, obese ferrets show decreased ability to induce antiviral responses and increased inflammatory signaling as assessed by gene ontology. Obese ferrets had decreased adiponectin (ADIPOQ) levels, similar to obese humans (*28*, *29*).

**Fig. 1.**
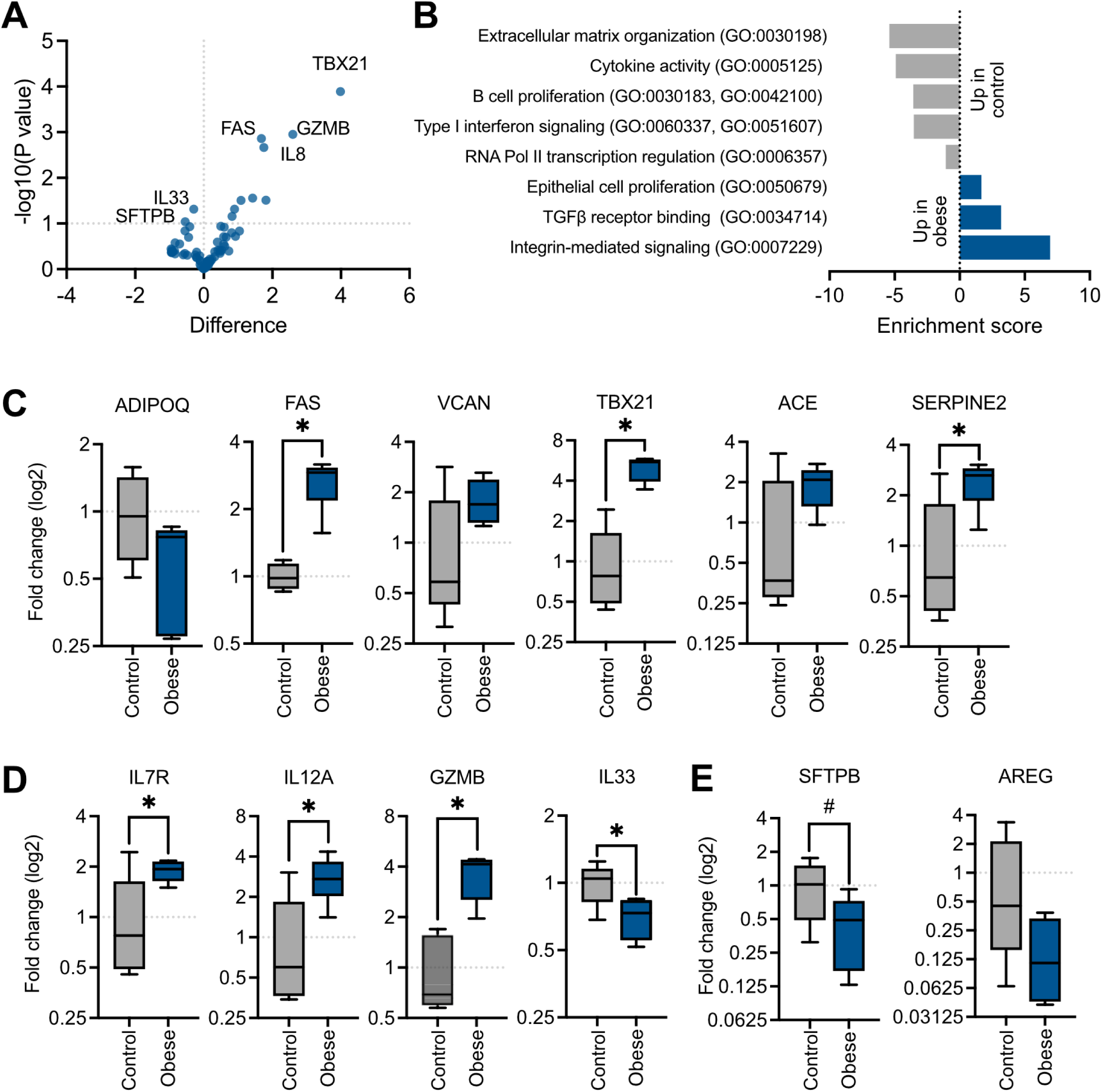
Obesogenic diet alters the pulmonary microenvironment at baseline. After 12 weeks on diet, obese and control ferrets were sacrificed and individual lobes of the lungs resected. Tissues were homogenized and RNA was extracted for high-throughput PCR. Each lobe was considered a separate sample. (A) Volcano plot of differences in gene expression between control and obese ferrets, represented by –log10 of *p*-value against log2-fold change. (B) DAVID enrichment analysis of up and down-regulated genes in uninfected obese ferret lungs. Absolute value indicates enrichment score, with negative scores indicating relative down-regulation from uninfected control lungs. (C-E) Differences in pulmonary gene expression between control and obese ferrets at homeostasis. Fold expression was calculated via the –ΔΔCt method. (C) Obesity-related markers are increased in obese ferrets. Expression of Fas cell surface death receptor (FAS, *p*=0.0014), T-box transcription factor 21 (TBX21), *p*=0.0001), and serpin family E member 2 (SERPINE2, *p*=0.0277) were significantly increased. (D) Obese ferrets have significantly increased levels of the immune and antiviral defense genes interleukin 17 receptor (IL7R, *p*=0.0484), interleukin 12A (IL12A, *p*=0.0310), granzyme B (GZMB, *p*=0.0011), and interleukin 33 (IL33, *p*=0.0484). (E) Decreased pulmonary surfactant (SFTPB, *p*=0.0911) and amphiregulin (AREG) in obese ferrets. Significance was determined by unpaired *t* test. Data represents the average of 5 lung lobes per n=1 ferret/group, one independent experiment of n=1/group. Samples were run in duplicate. Error bars represent minimum value to maximum value. \**p*<0.05, ^#^*p*<0.10.

Fas cell surface death receptor (FAS), versican (VCAN), T-box transcription factor 21 (TBX21), angiotensin I converting enzyme (ACE), and serpin family E member 2 (SERPINE2), metabolic markers associated with obesity (*30–35*), were increased in the obese group. In addition, the baseline pulmonary environment was pro-inflammatory. Elevated IL7R, interleukin 12A (IL12A), and granzyme B (GZMB) suggest the presence of natural killer (NK) cells and cytotoxic T cells (*36–38*). Conversely, expression of the alarmin protein IL33 was decreased in obese ferrets (*39*). We also noted reduced levels of surfactant protein B (SFTPB), a protein produced by epithelial cells involved in maintenance of surface tension in the lung (*40*), as well as decreased amphiregulin expression (AREG), suggesting impaired growth and proliferation of epithelial cells (*41*, *42*). This data suggests uninfected obese ferrets have impaired lung function and a pro-inflammatory environment even before infection, potentially altering viral pathogenesis in obese ferrets during influenza infection.

### Influenza disease severity is increased in obese ferrets

The 2009 H1N1 influenza pandemic identified obesity as a risk factor for poor disease outcome (*43*, *44*). We hypothesized that like humans, obese ferrets would experience increased disease severity during influenza infection. To test, control and obese ferrets were inoculated with 10^6^ TCID_50_ of A/California/04/2009 (H1N1) virus and we monitored weight, body temperature, and clinical signs for 12 days post-infection (dpi). Obese ferrets lost significantly more weight and had higher body temperature during the acute phase of infection (Fig. 2A-B). We observed more disease symptoms in the obese ferrets, such as lethargy, coughing, and nasal discharge (Fig. 2C).

**Fig. 2.**
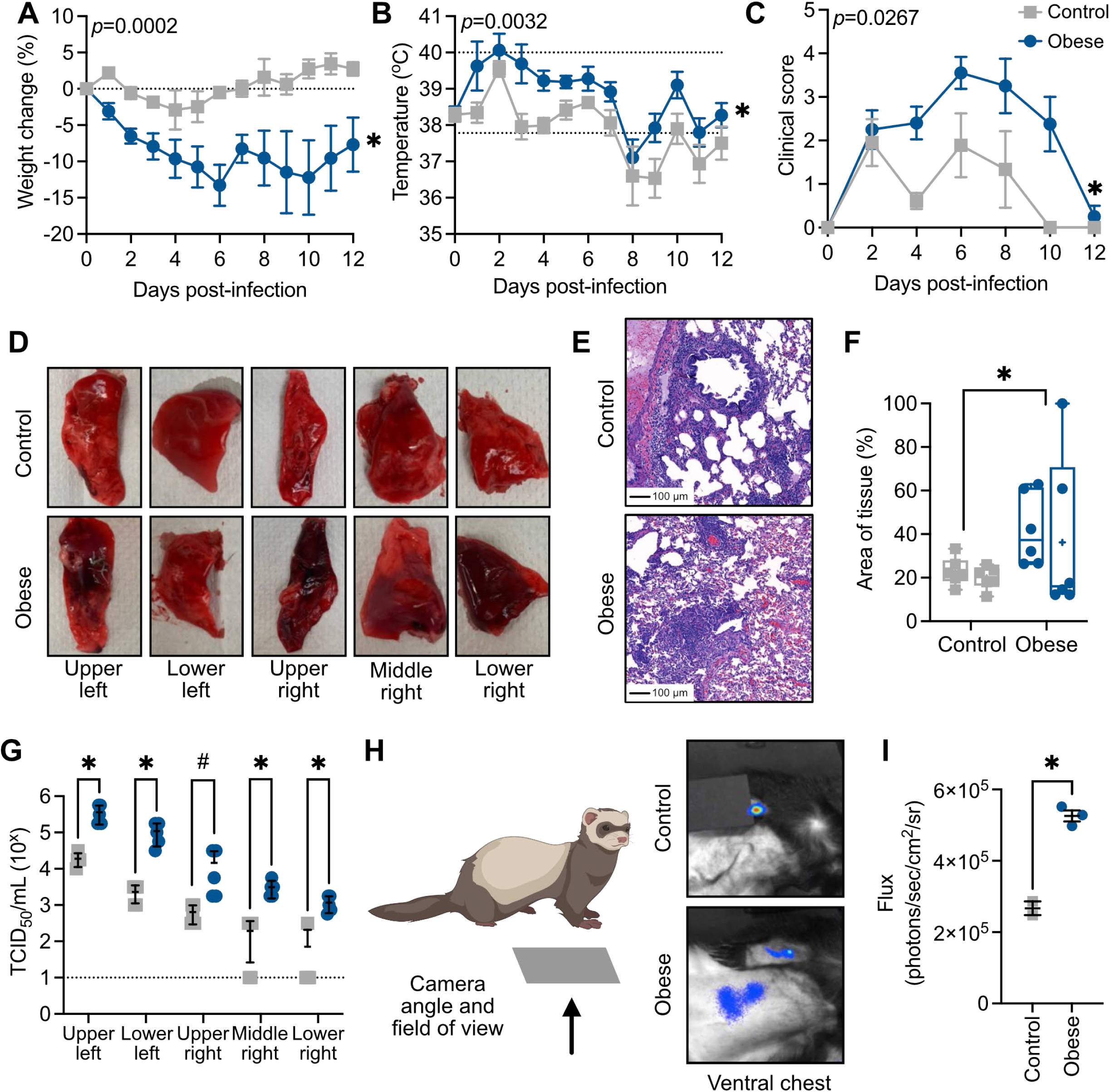
Disease severity following influenza infection is increased in obese ferrets. Control and obese ferrets were inoculated with 10^6^ TCID_50_ A/California/04/2009 (H1N1) virus and monitored for 12 days. (A) Weight (*p*=0.0002) and (B) body temperature (*p*=0.0032) were measured daily. (C) Clinical scores (*p*=0.0267) were recorded daily as described in methods. Data represents 2 independent experiments of n=4-6/group. (D) Gross pathology of lungs taken from control and obese ferrets infected with 10^6^ TCID_50_ A/California/04/2009 (H1N1) virus at 6 dpi. Images are representative of n=5/group. (E) Hematoxylin and eosin staining of lungs taken from control and obese ferrets infected with 10^6^ TCID_50_ A/California/04/2009 (H1N1) virus at 6 dpi. (F) Quantification of total inflammatory and immune infiltration area (*p*=0.0385) of (E). Data represents 1 independent experiment of n=2/group with 6 histology sections per ferret. (G) Lungs were separated by individual lobe and TCID_50_ assays performed to measure infectious viral titers (upper left lobe, *p*=0.0043; lower left, *p*=0.0079; upper right, *p*=0.0845; middle right, *p*=0.0034; and lower right, *p*=0.0031). Data represents 1 independent experiment with n=5/group. (H) Control and obese ferrets were infected with 10^5.5^ TCID_50_ of the A/California/04/2009-PA NLuc (H1N1) bioluminescent reporter virus. At 2 (*p*=0.0175) dpi, the chest was imaged to detect replicating virus in the lungs. Luminescence on the arm indicates the substrate injection site. (I) Quantification of (H). Data represents 1 independent experiment of n=3/group. Statistical significance was determined by mixed-effects analysis with repeated measures (A-C, *p*-value represents simple main effects analysis on diet), multiple unpaired t-tests (G), or nested t-test between independent ferrets and across diets (F). Error bars show standard error of the mean (SEM, A-C, G) or standard deviation of the mean (SD), (F, I). Dashed lines indicate baseline weight (A), normal range (B) or lower limit of detection (G). \**p*<0.05, ^#^*p*<0.10.

The increased clinical symptoms and disease severity led us to question whether there was greater progression to the lower respiratory tract (LRT) than in the control group. We infected male ferrets with A/California/04/2009 (H1N1) virus and collected lungs at 6 dpi. We noted increased damage in the lungs of the obese ferrets by gross pathology (Fig. 2D). Histological analysis showed increased inflammation and immune infiltration in obese compared to control lungs at 6 dpi (Fig. 2E-F), supporting an increase in overall disease severity in obese ferrets upon influenza virus infection. Viral titers were measured by TCID_50_ assay, treating the individual lobes of the lung as separate samples to spatially assess viral spread. Obese ferrets had overall significantly higher viral titers, even in the distal lung, suggesting that viral spread was increased (Fig. 2G) and/or faster viral clearance in the control group. To confirm more extensive viral spread at this timepoint, we infected control and obese ferrets with 10^5.5^ TCID_50_ of A/California/04/2009-PA NLuc, a bioluminescent reporter virus that allows longitudinal imaging of influenza virus (*45*, *46*). At 2 dpi, we detected significantly more luminescence in the obese ferrets including areas of the lower lung (Fig. 2H-I).

Of note, there were minimal differences in disease symptoms in female ferrets. Obese female ferrets were not significantly larger than the control group, and there were no significant differences in markers of metabolic syndrome (Fig. S2A-J). Disease severity was not significantly increased in obese female ferrets, although body temperature and clinical scores trended higher in the obese group (Fig. S2K-M).

### Obesity alters the pulmonary response to influenza infection

Next, we assessed the pulmonary response to infection in the context of obesity (Figure 3A). The gene expression profile at 6 dpi was chosen to assess lung repair and remodeling in addition to the transition from innate to adaptive immunity. Additionally, obese ferrets still had high viral titers throughout the lungs at this timepoint. First, we compared gene expression in obese and control lungs post-infection to control lungs at baseline to quantify global infection-related changes in gene expression. Interestingly, the gene profile of obese lungs at 6 dpi was more similar to uninfected lungs of both control and obese ferrets than to infected control lungs, indicating less of an overall change in antiviral state post-infection in the obese ferrets (Fig. 3A). Next, we determined how the expression of individual genes was changing in infected control and obese ferrets compared to levels in diet-matched uninfected lungs. Notably, key inflammatory mediators, such as interleukin 1 (IL1A) and nitric oxide synthase 2 (NOS2) were upregulated post-infection in obese ferrets, possibly leading to extensive inflammatory damage of the respiratory epithelium as observed in histopathology (Fig. 3B) (*47*, *48*). Conversely, genes involved in the innate interferon (IFN) response were downregulated in obese ferrets (Fig. 3C), as well as other antiviral defense mediators such as interferon-ε (IFNE), interferon-γ (IFNG), TBX21 and GZMB (Fig. 3D) suggesting dysregulation of adaptive immune cells (*49*, *50*). The antiviral mediator MX1 was lower in obese compared to control ferrets, mirroring results from both murine and human studies (Fig. 3C) (*51*, *52*). While key players in the inflammatory response, interleukin-6 (IL6.C) and interleukin-1b (IL1B) also have protective effects post-influenza infection (*53*, *54*) and show reduced upregulation in obese lungs compared to control. This could be due to the heightened pro-inflammatory environment already present in obese lungs at baseline compared to control lungs and contribute the dysregulated antiviral response in obese ferrets post-infection.

**Fig. 3.**
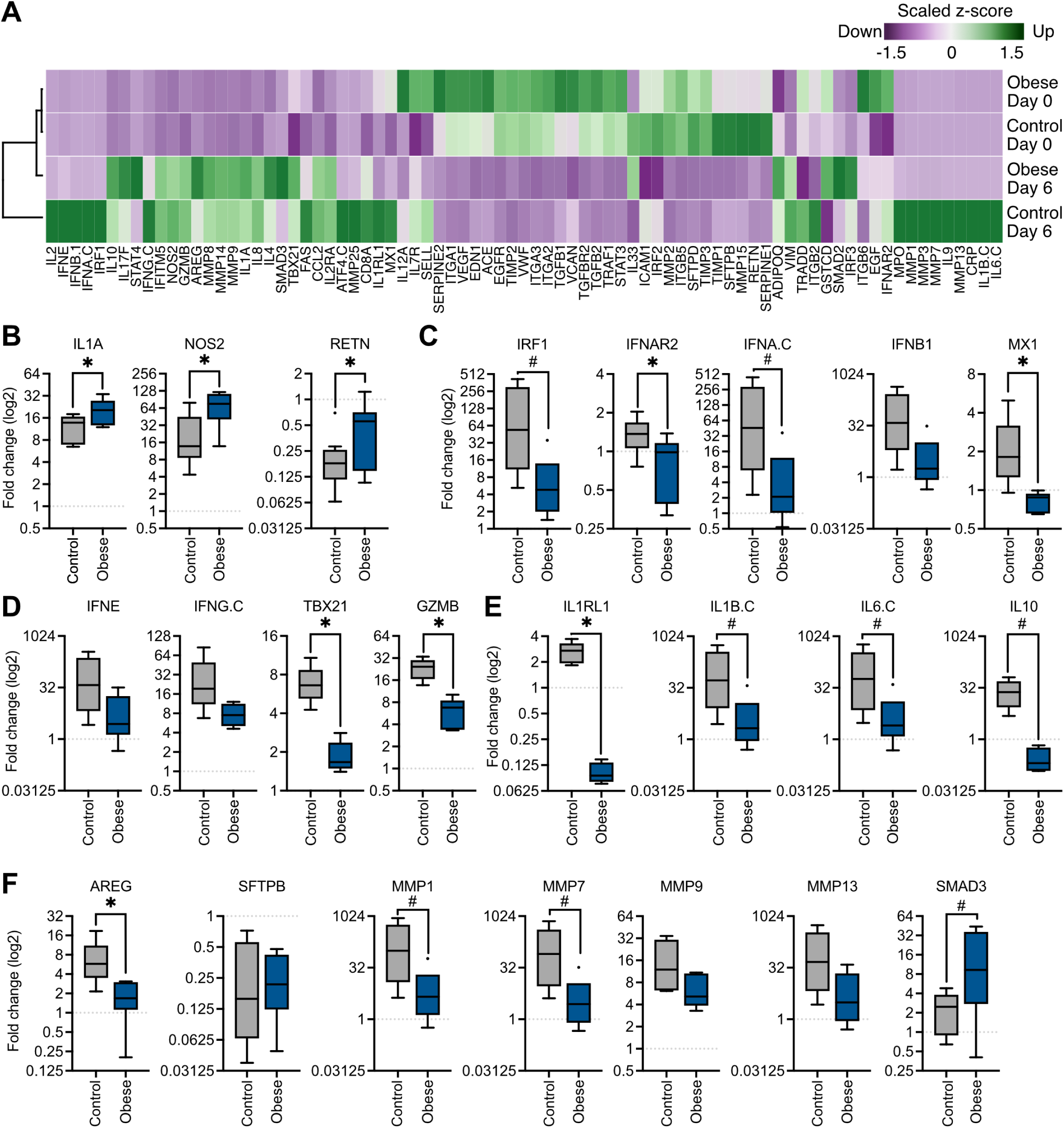
Altered inflammatory and immune gene expression in obese ferrets post-infection. Ferrets infected with 10^6^ TCID_50_ of A/California/04/2009 (H1N1) virus were sacrificed at 6 dpi to collect lung tissue. Lungs were separated by lobe and each lobe was treated as an individual sample. RNA was extracted from homogenates for high-throughput PCR. (A) Heatmap of gene expression normalized to uninfected control ferret using the 2^-ddCt^ method. Obese infected ferrets group more closely with obese baseline (bars). Green indicates increased gene expression relative to uninfected control ferret, and purple indicates decreased. (B-F) Gene expression in obese and control ferret lungs post-infection, normalized within diet groups to expression levels at baseline. (B) IL1A, *p*=0.0342; NOS2, *p*=0.0192; RETN, *p*=0.0449. (C) IRF1, *p*=0.0733; IFNAR2, *p*=0.0218; IFNA.C, *p*=0.0877; MX1, *p*=0.0129. (D) TBX21, *p*=0.0001; GZMB, *p*<0.0001. (E) IL1RL1, *p*=0.0003; IL1B.C, *p*=0.0934; IL6.C, *p*=0.0953; IL10, *p*=0.0535. (F) AREG, *p*=0.0110; MMP1, *p*=0.0833; MMP7, *p*=0.0890, SMAD3, *p*=0.0513. Significance was determined by unpaired *t* test. Data represents the average of 5 lung lobes per n=2 ferrets/group, one independent experiment of n=2/group. Samples were run in duplicate. Error bars represent minimum value to maximum value. \**p*<0.05, ^#^*p*<0.10.

In addition to this reduced innate response, obese ferrets had lower interferon-γ (IFNG) and interleukin 2 (IL2) expression, suggesting dysregulation of adaptive immune cells (*49*, *50*). Both interleukin 10 (IL10), a regulatory cytokine associated with decreased immunopathology during influenza infection (*55*), and interleukin 1 receptor like 1 (IL1RL1), the receptor for the alarmin IL33 that alerts of epithelial trauma (*39*), were downregulated in obese ferrets compared to control.

Finally, we quantitated genes involved in lung function and repair (Fig. 3F). SFTPB levels were reduced in both control and obese ferrets, possibly because influenza infection results in the death of epithelial cells (*56*). However, control ferrets upregulated AREG expression more than obese, suggesting an increase in epithelial cell regeneration (*41*, *42*), compared to obese. We also noted evidence of extracellular matrix remodeling in control ferrets as increased levels of matrix metalloproteases were detected post-infection. This data suggests obese ferrets may not adequately mount an immune response to infection and inefficiently repair lung tissue, however further investigation throughout the course of infection is needed. Future tool development is needed to validate these findings at the protein level.

### Increased disease severity in obese ferrets is not strain-specific

Human obesity is more strongly associated with increased risk for some influenza strains versus others (*6*, *8*, *57*). To determine how obesity impacts disease severity of different influenza virus strains, we chose representative viruses A/Memphis/257/2019 (human seasonal H3N2), A/Hong Kong/1073/1999 (avian-like H9N2), and B/Brisbane/60/2008 (influenza B, Victoria lineage) and challenged control and obese ferrets with 10^6^ TCID_50_ of H3N2 or H9N2, or 10^5.5^ TCID_50_ of influenza B viruses. During H3N2 and influenza B infections, obese ferrets lost more weight. Yet, there were no observable differences with H9N2 infection (Fig. 4A-C). Differences in body temperature were minimal (Fig. 4D-F) but the obese group trended slightly higher at 3-5 dpi during influenza B infection (Fig. 4F). Differences in disease severity were predominantly symptomatic. Obese ferrets had significantly increased clinical scores compared to control, notably nasal discharge, cough, and excessive sneezing (Fig. 4G-I). H9N2 infection was particularly interesting in that it caused no discernable symptoms in control ferrets yet obese ferrets had relatively high clinical scores. Further, we detected virus in the lungs of 2 H9N2-infected obese ferrets at 4 dpi; one in the lower left lobe (titer=10^4.75^ TCID50/mL) and the other the lower right (titer=10^4.75^ TCID50/mL). Taken together, obese ferrets were significantly more likely to experience severe symptoms during influenza infection independent of strain (Table 1).

**Fig. 4.**
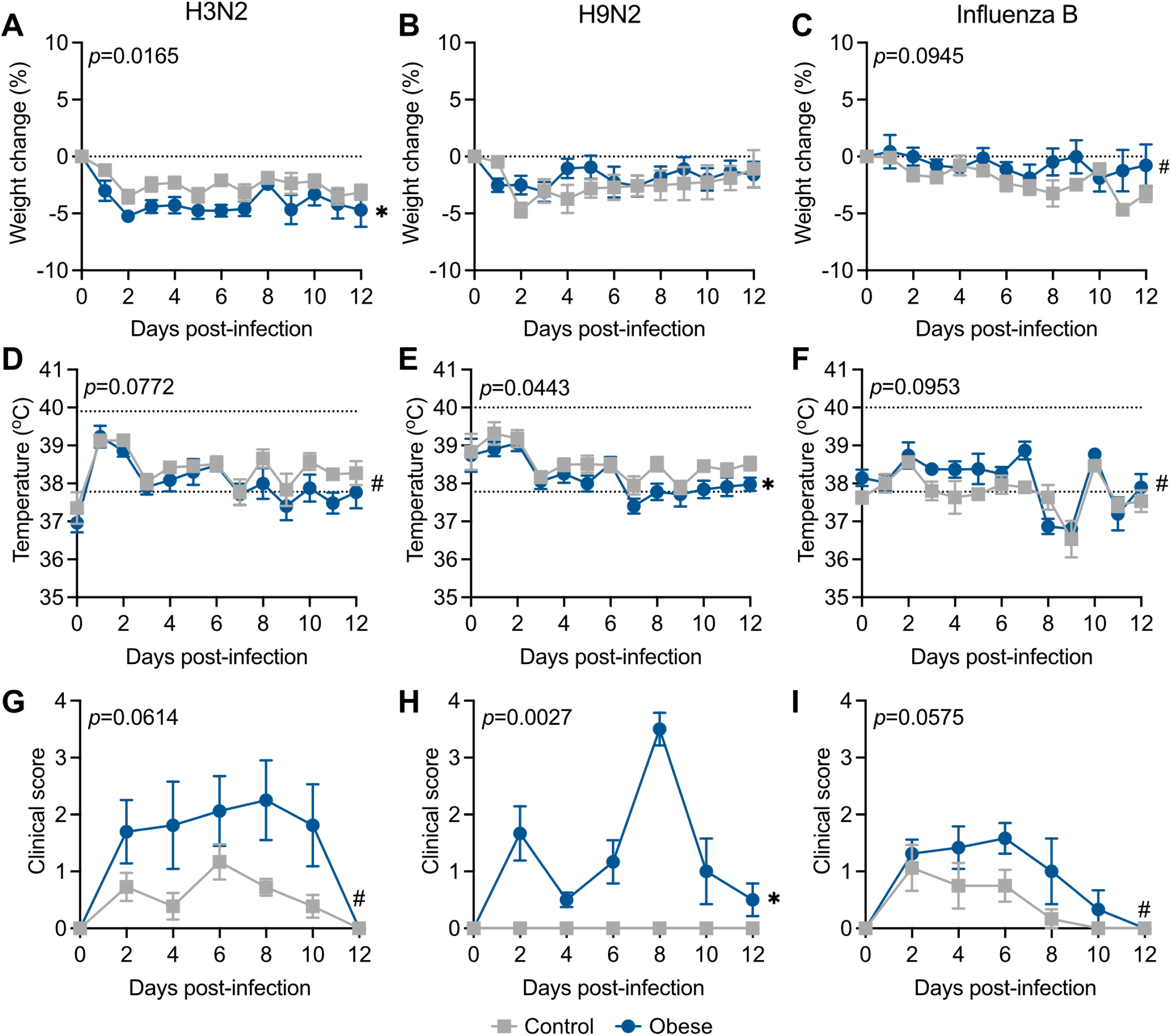
Increased disease severity in obese ferrets is independent of viral strain. Control and obese ferrets were infected with 10^6^ TCID_50_ of A/Memphis/257/2019 (H3N2), A/Hong Kong/1073/1999 (H9N2), or B/Brisbane/60/2008 (influenza B) viruses and monitored for 12 days. (A) Weights (H3N2 *p*=0.0165; H9N2 *p*=0.3492; influenza B *p*=0.0945), (B) body temperature (H3N2 *p*=0.0772; H9N2 *p*=0.0443; influenza B *p*=0.0953), (C) clinical scores (H3N2, *p*=0.0614; H9N2, *p*=0.0027; influenza B, *p*=0.0575), and (D) nasal wash scores (H3N2, *p*=0.0003; H9N2, *p*=0.0267; influenza B *p*=0.1953) were recorded as described in Figure 1. Data represents 2 independent experiments of n=6-7/group (H3N2), 4 independent experiments of n=2-7/group (H9N2), and 2 independent experiments of n=3-5/group (influenza B). Significance was determined by mixed-effects analysis with repeated measures (*p*-value represents simple main effects analysis of diet). Error bars represent standard error of the mean (SEM) and dashed lines indicate baseline weight (A-C) or normal range (D-F). \**p*<0.05, ^#^*p*<0.10.

**Table 1.** Risk of symptoms by diet and influenza strain.

| **Virus** | **Diet** | **Total ferrets** | **At-risk days** | **Symptomatic days** | **Average symptomatic days per ferret** | **Absolute risk** | **Relative risk for obese** | ***p*-value** |
| --- | --- | --- | --- | --- | --- | --- | --- | --- |
| **H1N1\*** | Control | 8 | 112 | 18 | 2.3 | 0.16 | 2.62±1.65 | <0.0001 |
|  | Obese | 8 | 88 | 37 | 4.9 | 0.42 |  |  |
| **H3N2†** | Control | 16 | 184 | 48 | 3.2 | 0.26 | 2.27±0.70 | 0.0001 |
|  | Obese | 17 | 198 | 117 | 7.0 | 0.59 |  |  |
| **H9N2‡** | Control | 13 | 146 | 15 | 1.1 | 0.10 | 1.91±1.57 | 0.03 |
|  | Obese | 13 | 122 | 24 | 1.7 | 0.20 |  |  |
| **Influenza B§** | Control | 8 | 68 | 16 | 1.9 | 0.24 | 1.75±1.18 | 0.03 |
|  | Obese | 8 | 68 | 28 | 3.5 | 0.41 |  |  |
| **Total** | Control | 45 | 510 | 97 | 2.1 | 0.19 | 2.28±0.52 | <0.0001 |
|  | Obese | 46 | 476 | 206 | 4.3 | 0.43 |  |  |

\*A/California/04/09; †A/Memphis/257/2019; ‡A/Hong Kong/1073/1999; §B/Brisbane/60/2008

### Obesity impacts viral replication in the upper respiratory tract

Because the upper respiratory tract (URT) is the primary site of infection, we next asked how obesity would impact viral replication in the nasal cavity by quantitating viral titers in nasal washes collected every 48 h during the experiments shown in Figs. 1 and 3.

Overall, obese ferrets trended towards increased viral titers in the URT, and in the case of H1N1 and H9N2 viruses, had detectable virus in the nasal washes for a longer duration (Fig. 5A). Differences in viral load were most pronounced by 4 dpi except for H9N2 virus, where viral replication was increased as early as 2 dpi in the obese group. Viral replication at the peak of infection in the obese ferrets was higher during H1N1, H3N2, and H9N2 infection but not influenza B. We could visualize significant increases in luminescent flux in the nasal cavities of obese ferrets at 2 and 4 dpi using the bioluminescent reporter virus A/California/04/2009-PA NLuc (H1N1) as in Fig. 1, and signal in the obese ferrets persisted longer (Fig. 5B-C).

**Fig. 5.**
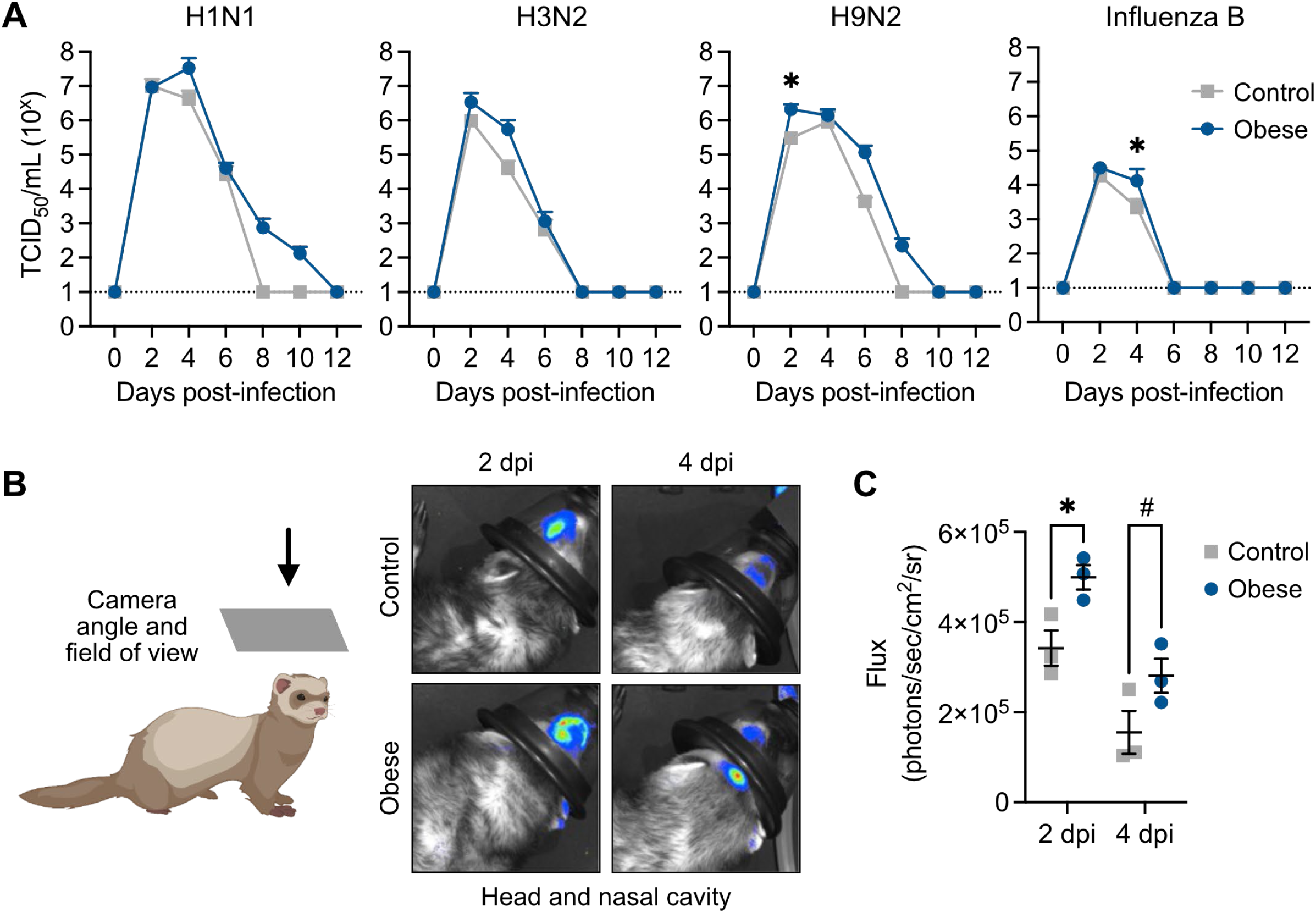
Increased viral replication in the upper respiratory tract of obese ferrets. (A) Control and obese ferrets were infected with 10^6^ TCID_50_ of A/California/04/2009 (H1N1), A/Memphis/257/2019 (H3N2), A/Hong Kong/1073/1999 (H9N2), or 10^5.5^ TCID_50_ of B/Brisbane/60/2008 (influenza B) viruses. Nasal washes were collected every 48 h and viral titers measured by TCID_50_ assay. H1N1 4 dpi *p*=0.2793; H9N2, 2 dpi *p*=0.0275 and influenza B, 2 dpi *p*=0.0476 by 2-way ANOVA with multiple comparisons. Data represents two independent experiments of n=4-7/group. (B) Control and obese ferrets were infected with 10^5.5^ TCID_50_ of A/California/04/2009-PA NLuc (H1N1) virus and the nasal cavity was imaged at 2 (*p*=0.0412) and 4 (*p*=0.0995) dpi. (C) Quantification of (B). Statistical significance was determined by 2-way ANOVA with multiple comparisons. Data represents 1 independent experiment of n=3/group. Error bars show standard error of the mean (SEM) (A) or standard deviation of the mean (SD) (D) and dashed lines indicate the lower limit of detection. \**p*<0.05, ^#^*p*<0.10.

### Obesity can impact viral transmission dynamics of a zoonotic virus

Due to the increased susceptibility of obese individuals to certain influenza strains (*6*), and the reduced antiviral response at 6 dpi observed in the obese ferrets (Fig. 3), we hypothesized obese ferrets might be more susceptible to influenza infection. Since circulating human influenza strains transmit easily in the ferret model, we chose an avian-like virus, A/Hong Kong/1073/1999 (H9N2), with documented limited transmission ability between ferrets (*58*). Briefly, control ferrets were inoculated with 10^6^ TCID_50_ of H9N2 virus and 24 hours later, a control or obese naïve contact ferret was introduced into the same cage as an infected index ferret. Each pair was in direct contact and could interact freely. Nasal washes were collected from both the index and contact ferrets every 48 h to monitor viral transmission, and at 21 dpi blood was collected to assess seroconversion. We included both homologous and heterologous diet pairings (Fig. 6A).

**Fig. 6.**
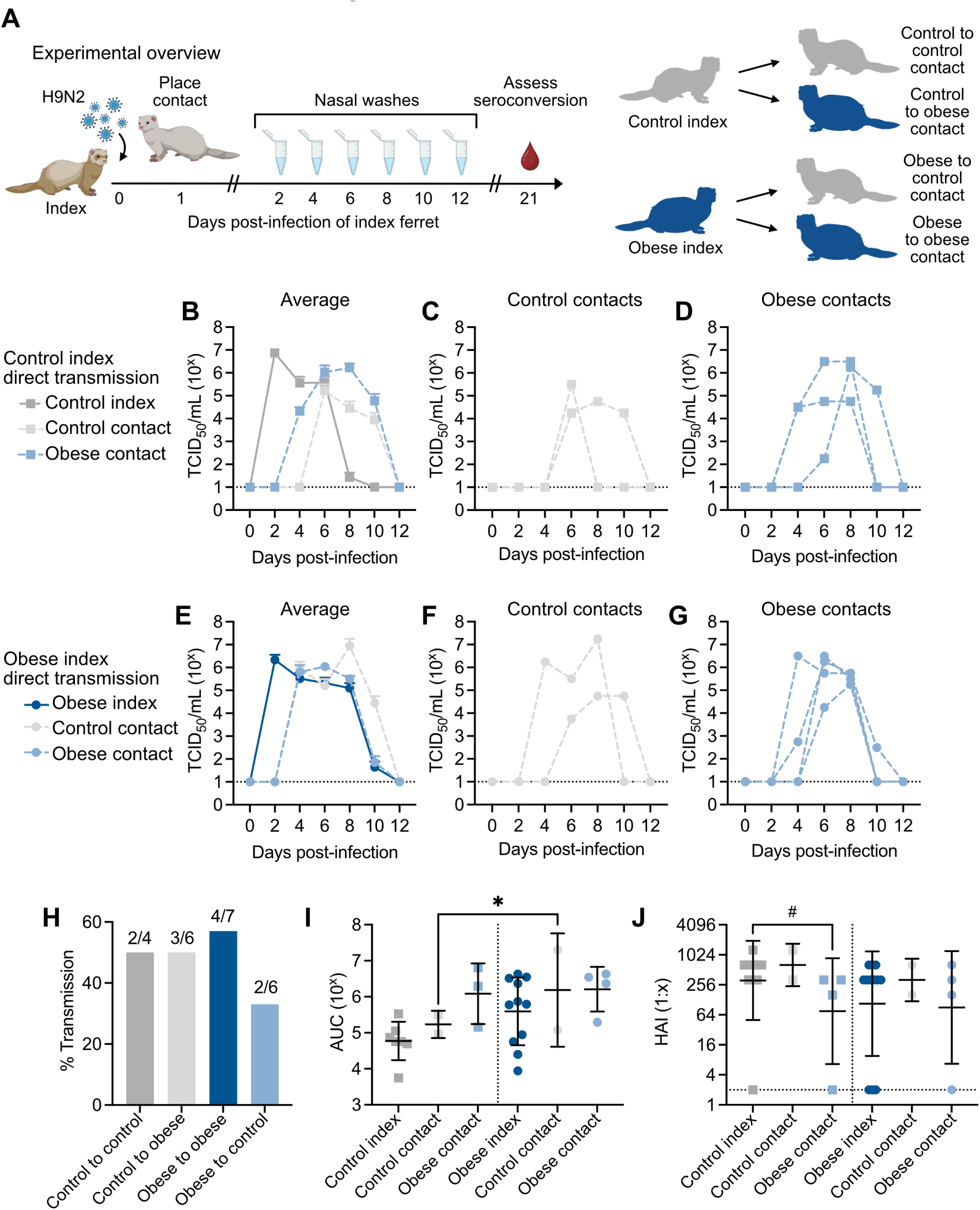
Obesity can impact viral transmission dynamics. (A) Experimental setup for a direct transmission study. An index ferret was inoculated with 10^6^ TCID_50_ A/Hong Kong/1073/1999 (H9N2) virus. After 24 h, a naïve contact ferret was introduced. Nasal washes were collected every 48 h for 12 days and blood was collected at 21 dpi. (B) Nasal wash titers of transmission groups with a control index ferret were measured by TCID_50_ assay. (C) Nasal wash titers of individual control and (D) obese contacts paired with control index. Number in top left indicates the number of ferrets that became influenza-positive/total number of contact ferrets. (E) Nasal wash titers of transmission groups with an obese index ferret. (F) Nasal wash titers of individual control and (G) obese contacts paired with obese. (H) Total number of influenza positive contact ferrets for each transmission scheme (I) Area under the curve (AUC) of (B) and (E), control contacts *p*=0.0270 by one way ANOVA with Sidak’s test. (J) Hemagglutination inhibition assays were performed on plasma collected at 21 dpi. Control index vs obese contact paired with control *p*=0.0793 by unpaired *t* test. Data represents 3 independent experiments of n=3-4/group. Error bars indicate standard error of the mean (SEM) and dashed lines indicate lower limit of detection in (B-G, J). \**p*<0.05, ^#^*p*<0.10.

When paired with control index ferrets, obese contacts retained the phenotype of higher viral titers and increased shed duration compared to control contacts (Fig. 6B-D, Table 2). Regardless of the diet status of the contact ferret, the transmission rate of H9N2 virus from a control index ferret was 50%; however, obese contacts became infected earlier and shed virus longer than control contacts (Fig. 6D, Table 2). We concluded that while transmission efficacy did not appear to be altered based on the diet of the contact ferret, the obese ferrets may be more susceptible to infection since 2 of 3 obese contacts became positive for influenza virus earlier than control contacts (Fig. 6D, Table 2).

**Table 2.**
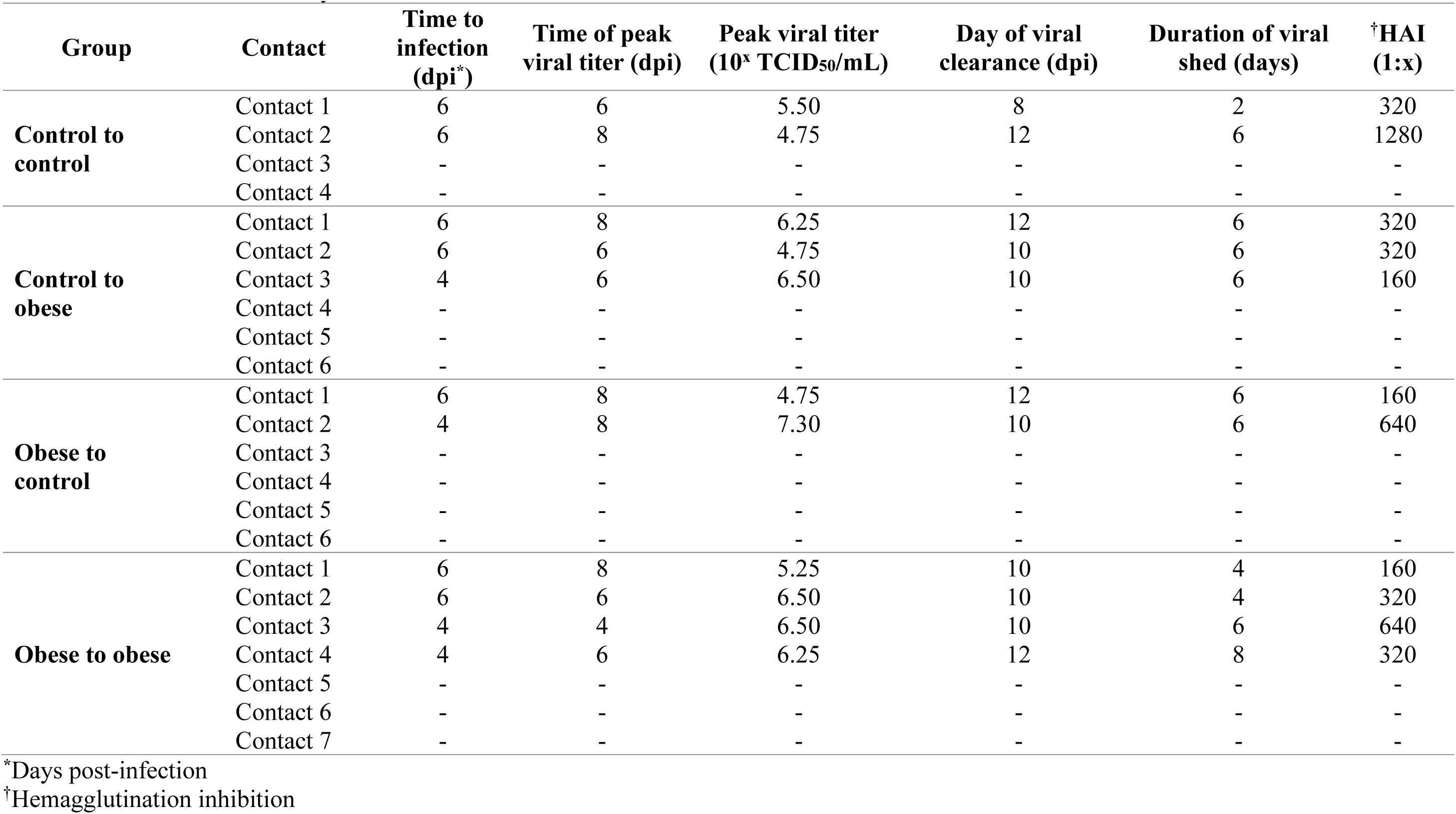
H9N2 transmission dynamics.

We next considered the possibility that obese ferrets may be more likely to transmit, as increased viral shed can facilitate transmission (*59*). We repeated the experiment using obese ferrets as indices. Again, obese contacts retained the phenotype of increased viral titers and duration of shed (Fig. 6E-G, Table 2). Obese ferrets transmitted to only 2 of 6 control contacts (33%, Fig. 6H), with only one control contact displaying robust viral shed. Conversely, obese ferrets successfully transmitted to 4 of 7 obese contacts (57%, Fig. 6H). The viral titers of obese ferrets had similar area under the curve (AUC) regardless of index diet. While we detected active viral replication in only 2 control contact ferrets making statistical comparisons difficult due to low numbers and high variation, we did note greater AUC in control ferrets paired with obese index compared to those paired with control (Fig. 6I). Intriguingly, obese contacts shed greater quantities and for longer than control index ferrets, while no difference in AUC was noted in obese index pairings (Fig. 6I). Finally, when assessing seroconversion by hemagglutination inhibition (HAI) assay, obese contacts paired with control ferrets had decreased HAI titers compared to control contacts paired with control index. Further, control contacts paired with obese index trended lower that control contacts paired with control (Fig. 6J, Table 2). Taken together, obesity may influence transmission dynamics of H9N2 virus.

## Discussion

In 2022, approximately 800 million adults, children, and adolescents were considered obese (BMI>30). In some countries, over 40% of the population are obese with levels continuing to rise. This is concerning given the health risks associated with a high BMI including increased risk of developing severe disease with respiratory viruses like SARS-CoV-2 and influenza. In these studies, we developed an obese ferret model to study the impact of obesity on disease severity and transmission of influenza virus. Like humans (*6*), obese ferrets experienced increased disease severity during H1N1 infection as shown by increased weight loss and body temperature, as well as greater severity and duration of symptoms. Viral spread and lung injury were increased in obese ferrets, which is similar to humans where case reports of bronchiolitis, alveolar hemorrhage, and increased virus in the lungs of obese decedents has been reported (*60*). As ferrets are known to recapitulate the human respiratory tract, our model helps translate these discoveries to the clinic. While rodent models have provided valuable insight into mechanisms of disease severity in the context of obesity (*14*, *61*), the obese ferret and associated reagents provide a robust and translatable model organism for vaccine and antiviral testing and for risk assessments in surveillance and epidemic response scenarios.

In our model, we employed influenza strains displaying a spectrum of symptom severity from mild to severe. For each virus we tested, the obese group was more likely to experience more severe and prolonged symptoms compared to control (Table 1). Surprisingly, H9N2, a virus typically not detected in the lungs of ferrets (*62*), spread to the lower respiratory tract of 2 of the obese ferrets yet caused no observable clinical symptoms in control ferrets (Fig. 4). Additionally, obese ferrets had higher viral titers in the nasal wash, congruent with data suggesting obese individuals experience increased viral shedding for longer periods of time during both H1N1 and H3N2 infection compared to non-obese, even when asymptomatic (*7*). In our hands, the obese ferrets experienced prolonged viral shed in the case of H1N1 and H9N2 infections but cleared virus on the same day as control ferrets during H3N2 and influenza B infections. Human cohort studies also report strain-specific phenomena; one study found obesity had no bearing on human symptom severity during seasonal H3N2 infection as opposed to H1N1 (*6*), and indeed obese ferrets had lower symptom scores during H3N2 infection compared to H1N1.

The increased viral spread to the lower respiratory tract in obese ferrets could be explained by the dampened antiviral response in the lungs of obese ferrets. Blunted type I interferon responses are described in both mouse models and human epithelial cells *in vitro* (*17*, *18*). Further, obese ferrets did not adequately upregulate amphiregulin, MMPs, or the alarmin-activating receptor IL1RLI, genes involved in epithelial regeneration and wound healing. Obese ferrets also had lower levels of surfactant protein B and amphiregulin at baseline, which could compromise tensile strength of the lung and maintenance and turnover of the pulmonary epithelium leaving the lungs in an inherently weakened state. Additionally, homeostasis in the obese ferrets was characterized by high levels of inflammatory cytokines and pro-apoptotic protein FAS, which could compromise lung function.

Obesity also impacted viral transmission of an avian-like influenza virus. Obese ferrets were more susceptible to infection through direct contact independently of the index ferret’s diet. Although the rate of transmission from a control index was the same whether the contact was control or obese (50%), the obese ferrets were likely to become infected earlier. Unusually, an avian influenza virus transmitted more efficiently from obese index ferrets to obese contacts (57%), suggesting obesity impacts both transmission efficiency and susceptibility in the ferret model. Despite the lower likelihood of transmission from an obese to a control ferret (33%), one of the 2 control contacts that became infected shed virus earlier, had higher viral loads, and prolonged shed similar to our findings in obese ferrets. Honce *et al* reported that influenza viruses became more virulent as they were passed through an obese host but not control (*18*), suggesting that in some cases, virus shed by the obese ferrets may cause more severe infection. Using the more translatable ferret model to characterize the impact of obesity on viral evolution and bottlenecks in transmission dynamics is warranted as the rate of obesity continues to rise and may be valuable as a tool in risk assessment of emerging influenza viral strains.

Interestingly, increased disease severity during H1N1 infection could not be replicated using female ferrets. Metabolism can be sex-dependent (*63*), and this may explain the less drastic difference in disease severity in females; however, all ferrets used in this study were neutered or ovariectomized. In humans, different hallmarks of metabolic syndrome are also more likely to occur in males versus females (*64*). Metabolic dysfunction can also occur in the absence of phenotypic adiposity, leading to consequences such as diabetes and cardiovascular disease; conversely, individuals with larger body mass indices (BMI) can be metabolically healthy (*65*, *66*). In female ferrets, our obese diet protocol did not significantly increase ferret body mass or alter biochemical hallmarks of metabolic dysfunction, suggesting a different or prolonged approach may be necessary to induce ferret obesity and recapitulate increased disease severity in female ferrets. Further, applying our model to intact ferrets, or in other high-risk presentations such as pregnancy or age status, could shed light on the interactions among sex hormones, metabolic dysfunction, and immune responses on influenza disease outcomes (reviewed in (*67*)).

With every study, any caveats and limitations must be considered. Our study and those involving human cohorts were conducted with different methodology. For example, we defined viral shed by the detection of infectious virus, while in human studies the presence of viral genome is typically detected by the more sensitive qPCR assay (*6*, *7*, *59*). Further, the definition of symptom severity in clinical studies varies, and is often self-reported data that includes criteria non-applicable to ferret studies, such as hospitalization, ICU admission, and death. Observational scoring is also a subjective metric, so we used multiple persons to independently score our ferrets. Although we assessed gene expression in the ferret lung, due to the paucity of ferret reagents, we were unable to validate these findings at the protein level. Optimization of our primers is ongoing as improved genome builds become public. Finally, ferret studies are typically low *n* due to complex husbandry needs, limiting our statistical power (*12*). However, our studies align with and further a large body of research spanning work in cell culture, mouse models, immunology and epidemiology and point to a conserved effect of metabolic dysfunction on the viral infection dynamics.

Here we show ferrets fed calorically dense diets recapitulate features of obesity and metabolic dysfunction seen in humans and display increased influenza-induced disease severity. The new ferret reagents created for this study help us unravel the mechanisms of increased viral pathogenesis associated with obesity, including reduced antiviral responses and impaired wound healing, and will be key for studies beyond investigating the impact of diet on infectious disease. We report obesity influences both transmission dynamics and infection kinetics, potentially due to the increased viral replication in the URT and the prolonged duration of viral shed. Beyond obesity, future studies using our model could be used to determine what aspects of metabolic syndrome most effect influenza and other disease outcomes. Further, this model will be invaluable for testing next-generation vaccine and antiviral designs, as these pharmaceutical interventions traditionally show poor protection in durability in obese populations (*68*, *69*).These results, together with the increasing availability of ferret reagents (*70*, *71*), can help us understand many aspects of influenza disease in the context of obesity and beyond, such as immunological responses, risk assessment, and refinement of vaccines, antivirals, and other therapeutics to help protect this and other high-risk populations.

## Materials and Methods

### Experimental design

The objective of the study was to determine if we could determine mechanisms of disease severity in the context of obesity using the gold-standard animal model for influenza studies. Pre-study, ferret diets were optimized with the supervision of veterinarians in the St. Jude Children’s Research Hospital Animal Resource Center, and primers targeting genes of interest were stringently validated.

### Ethics statement

All animal experiments (protocol #513) were approved by the St. Jude Children’s Research Hospital Institutional Animal Care and Use Committee (IACUC). Ferrets were housed at ambient temperature (20°C and 45% relative humidity) with 12-hour light cycles and provided species and study appropriate enrichment activities.

### Ferret diets and measurements

Neutered, de-scented male ferrets or ovariectomized female ferrets were obtained from Triple F Farms (Elmyra, NY) at the age of 6 weeks and randomly assigned to obese or control diet groups. Diets were allocated daily for 12 weeks (Table S1). All ferrets were provided water *ad libitum*. Control ferrets were provided a restricted volume of high-density ferret chow (Lab Supply 5LI4; carbohydrates 17%, protein 36%, fat 47%) once daily. Obese ferrets received a 1:1 mix of high-density ferret diet plus feline diet (Lab Supply 5003; carbohydrates 40%, protein 32%, fat 27%) *ad libitum* and supplemented with wet kitten food (Iams 131321, Chewy.com) once per day beginning at 9 weeks of age. Throughout the diet period, physical measurements were recorded weekly. Ferret body length was measured from nose to base of tail and waist circumference measures just above the iliac crest. Skinfold fat was determined using a digital fat caliper on the right side of the abdomen above the iliac crest. Ferret mass index (FMI) was calculated by standardizing the product of weight in kilograms and circumference in centimeters by the ferret length in centimeters squared.

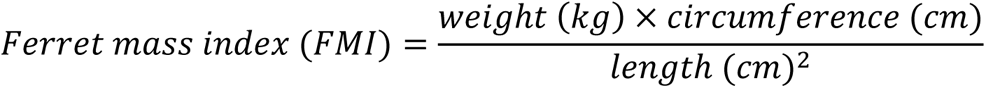

### Blood chemistry analyses

Ferrets were sedated with 4% isoflurane and blood was collected via the jugular vein into a heparinized collection tube (BD 366643). Plasma was separated via centrifugation and stored at –80°C until use. Blood chemistry analyses were performed by the Veterinary Pathology Core (VPC) at St. Jude Children’s Research Hospital. Leptin levels were determined using Human Leptin Instant ELISA kit (Invitrogen BMS2039INST) according to manufacturer’s instructions.

### Viruses and cells

Madin-Darby canine kidney (MDCK) cells (ATCC CCL-34) were cultured in Minimal Essential Medium (MEM) (Corning 10-010-CV) supplemented with 10% fetal bovine serum (FBS) (HyClone) and 200 mM L-glutamine (Gibco 35050079). Cells were incubated at 37°C/ 5% CO_2_.

To propagate A/California/04/2009 (H1N1), A/California/04/2009-PA NLuc (H1N1), A/Memphis/257/2019 (H3N2), and B/Brisbane/60/2008 (Victoria lineage) viruses, MDCK cells were inoculated with virus diluted in infection media [MEM supplemented with 200 mM L-glutamine, 0.075% bovine serum albumin (BSA) (Gibco 15260-037), and 1 μg/mL N-tosyl-L-phenylalanine chloromethyl ketone (TPCK)-treated trypsin (Worthington LS003740)] for 72 hours at 37°C/ 5% CO_2_. Supernatants were harvested, clarified by centrifugation, and aliquots stored at –80°C until use. A/Hong Kong/1073/1999 (H9N2) virus was propagated in 9-day old embryonated chicken eggs as previously described (*72*). Viral titers were measured by tissue culture infectious dose 50 (TCID_50_). Briefly, MDCK cells were seeded in 96-well plates at a density of 30,000 per well and inoculated in triplicate with tenfold serial dilutions of samples or viral stock diluted in infection media. Cells were incubated for 72 hours at 37°C/ 5% CO_2_. Viral titers were read by hemagglutination of 0.5% packed turkey red blood cells and calculated by the method of Reed and Muench (*73*).

### Infections

Ferrets were lightly sedated with 4% inhaled isoflurane and inoculated with virus diluted in phosphate-buffered saline (PBS) (Corning 21-040-CV) supplemented with 100 U/mL penicillin and 100 μg/mL streptomycin (Corning 30-002 CI). The dose of virus given was as follows: 10^6^ TCID_50_ of A/California/04/2009 (H1N1), A/Memphis/257/2019 (H3N2), A/Hong Kong/1073/1999 (H9N2); 10^5.5^ TCID_50_ of A/California/04/2009-PA NLuc (H1N1) and B/Brisbane/60/2008 (influenza B, Victoria lineage). Weight and temperature were assessed daily. Clinical scores were assessed by observing the presence and severity of symptoms using a point system. Metrics included sneezing (none=0, mild=1, excessive=2), coughing (absence=0, presence=1), nasal discharge (absence=0, presence=1), conjunctivitis (absence=0, presence=1), lethargy (active and playful=0, active when stimulated=1, not active when stimulated=2), and nasal wash consistency.

Mild sneezing was defined as 1-2 sneezes during the observation period, while continuous sneezing was considered excessive. Clinical scores were assessed by at least two investigators. Nasal washes were graded as follows: clear=0, cloudy=1, mucus present=2, mucus present with discoloration=3. Data is presented as the sum of points.

### Tissue collection

Ferrets were humanely euthanized via cardiac injection of Euthasol (Patterson Veterinary Supply). Animals sacrificed after 12 weeks on diet were necropsied. Abdominal adipose tissue was removed and weighed. For infection studies, individual lobes of the lungs were resected carefully to avoid cross-contamination. Lung tissue were finely minced with scissors before adding 1 mL sterile PBS and bead beaten. Samples were centrifuged for 5 min at 1,500 *g* to clarify before measuring viral titers by TCID_50_ assay. For histological analysis, whole tissues were submerged in 10% neutral buffered formalin for at least 48 hours, followed by tissue mounting, sectioning, and staining by the VPC at St. Jude Children’s Research Hospital.

### Bioluminescent imaging

Ferrets infected intranasally with A/California/04/2009-PA NLuc virus were imaged as previously described (*45*). Briefly, ferrets were anesthetized with 4% inhaled isoflurane and the chest area directly above the lung was shaved to minimize background. Ferrets were injected via the cephalic vein with Nano-Glo substrate (Promega N1110) at a dilution of 1:5 in sterile PBS. Ferrets were imaged using a Xenogen IVIS200 (PerkinElmer) with an exposure time of 5 min. Data was analyzed with LivingImage software (PerkinElmer).

### Nasal wash collection

Ferrets were anesthetized intramuscularly with 30 mg/kg ketamine (Patterson Veterinary Supply) and sneezing was induced by adding 1 mL PBS supplemented with 100 U/mL penicillin and 100 ug/mL streptomycin dropwise to the nares. Sample was collected into a sterile specimen cup, briefly centrifuged, and stored at –80°C until use. Viral titers were determined via TCID_50_ assay as described.

### High-throughput qPCR and primer validation

Primers targeting immune and lung function-related genes of interest were designed based on the ferret genome sequences provided by Ensembl (https://useast.ensembl.org/Mustela_putorius_furo/Info/Index). Primer validation was conducted as described by the manufacturer (*74*). Tissues were collected from multiple ferrets and RNA was extracted using the RNeasy RNA extraction kit (Qiagen 74104) including an on-column DNase digestion using the RNAse-Free DNase set (Qiagen 79254). RNA was quality-assessed and quantified using a NanoDrop 8000 spectrophotometer. cDNA was synthesized with the SuperScript IV VILO master mix (Invitrogen 11756050) per manufacturer’s instructions. Specific target amplification was performed with Preamplification Master Mix (Fluidigm PN 100-5580) and candidate primers targeting genes of interest (Table S2) at 95°C for 10 minutes, followed by 12 cycles of 95°C for 15 seconds and 60°C for 4 min followed by exonuclease I (New England Biolabs, PN M0293S) treatment at 37°C for 30 minutes and 80°C for 15 minutes to remove any unincorporated primers. The resulting samples were diluted 1:5 using DNA suspension buffer (Teknova T0221). These samples were considered the sample at the highest concentration. Using DNA suspension buffer, we made 15 step 2-fold dilution series in DNA suspension buffer. Samples were prepared with SsoFast EvaGreen Supermix (BioRad PN 172-5211), loaded onto a primed 96.96 internal fluidics chip (IFC) (Fluidigm BMK-96.96) and run in triplicate against each individual primer to be validated along with at least 3 non-template controls. Data was analyzed using a Fluidigm Biomark HD and Fluidigm Real Time PCR Analysis Version 4.7.1. Ct values that fell in the range of 5-24 for each combination of assays were accepted. ΔCt values for each dilution step from the previous dilution samples were calculated and used to plot the mean slope of ΔΔCt against template concentration. The ideal slope would be 0, and we considered validated assay to fall in the range of –0.1 to 0.1. To further validate, PCR product was run on an agarose gel, bands were extracted, and sequenced via the Sanger method to ensure the correct gene was amplified. After validation, experimental samples were prepared using the above protocol, however the preamplification step was done using pooled primers. Experimental samples were run in duplicate and analyzed via the –ΔΔCt method. Data were normalized to expression at either baseline within diet groups or baseline of control, uninfected lungs. Gene expression visualized using R function *heatmap* with normalized gene expression scaled using the *scale* function and heatmap clustered based on expression patterns in samples and genes (*75*, *76*). Up– and down-regulated gene ontology was completed using DAVID functional annotation clustering with high classification stringency in the *Homo sapiens* gene background.

### Transmission experiments

Singly housed ferrets were lightly sedated with 4% inhaled isoflurane and inoculated with indicated virus diluted in 500 μL PBS. Directly inoculated ferrets are referred to as index ferrets. After 24 hours, a naïve ferret was placed in the same cage with the index ferret.

Ferrets could interact freely. For transmission between unlike diet groups, diets remained fixed for the index ferret. Weight, body temperature, and clinical signs were monitored as before. Nasal washes were collected every 48 h to monitor viral transmission.

### Hemagglutination inhibition assay

Blood was collected as described. Samples were centrifuged at 5,000 *g* for 15 min to separate the plasma and stored at –80°C until use. The red blood cell pellet was discarded. Samples were incubated with receptor destroying enzyme (RDE; Denka Seiken, Tokyo) overnight at 37°C. RDE-treated samples were inactivated at 56°C for 1 h. PBS was added to a final dilution of 1:10 and samples were stored at –80°C for at least 4 h to ensure complete inactivation of any residual neuraminidase. Two-fold sample dilutions were added to a 96-well plate and incubated with 4 hemagglutinating units (HAU) of the indicated virus for 15 min at room temperature. Packed turkey red blood cells (0.5%) diluted in PBS were added to each well and plates were incubated at 4°C for 90 min. HAI titer is reported as the reciprocal dilution of the last positive well. Positive and negative controls, as well as back titrations of virus, were included on each individual plate. If back titration value was >1:8 the assay was repeated.

### Statistical analysis

Data was analyzed using GraphPad Prism version 9.5.1 using the appropriate statistical test as indicated in figure legends. Final figures were prepared using Affinity Designer (Serif).

## Acknowledgments

The authors gratefully acknowledge the St Jude Animal Resource Center and Dr. Ana Vazquez-Pagan, Bridgett Sharp, and Sean Cherry for their help with ferret husbandry and feeding. We thank Drs. Harshan Pisharath, Amy Funk and Tiffani Rogers, and the veterinary nutritionists at Lab Diets for their valuable help ensuring the ferret diets provided appropriate nutrition. We thank the St Jude Veterinary Pathology Core for their help with blood analytics. Illustrations in Figure 1, 4, and 5 created with BioRender.com.

## Obese ferret model

United States patent application PCT/US2021/041230 (VM, RH, SSC)

## Funding

This work was funded by:

National Institutes of Health Centers for Excellence in Influenza Research and Surveillance (CEIRS) contract HHSN27220140006C (SSC)

National Institutes of Health Centers for Excellence in Influenza Research and Response (CEIRR) contract 75N93021C00016 (SSC)

National Institutes of Health Collaborative Influenza Vaccine Innovation Centers (CIVIC) contract 75N93019C00052

National Institutes of Health grant R01AI140766 (SSC)

The American Lebanese Syrian Associated Charities (ALSAC) (SSC)

## Author contributions

Conceptualization: SSC, VM, RH, EK

Methodology: SSC, VM, RH, EKA, DB, PT

Investigation: VM, RH, BL, EKA, DB, EK, PF, LL, VH, HT, PHB

Visualization: RH, VM

Supervision: SSC, RH, VM

Writing – original draft: VM, RH, SSC

Writing – review and editing: VM, RH, SSC, PHB, VH, PF, EKA

## Data and materials availability

All data are available in the main text or the supplementary materials.

## Supplementary Materials for

**Fig. S1.**
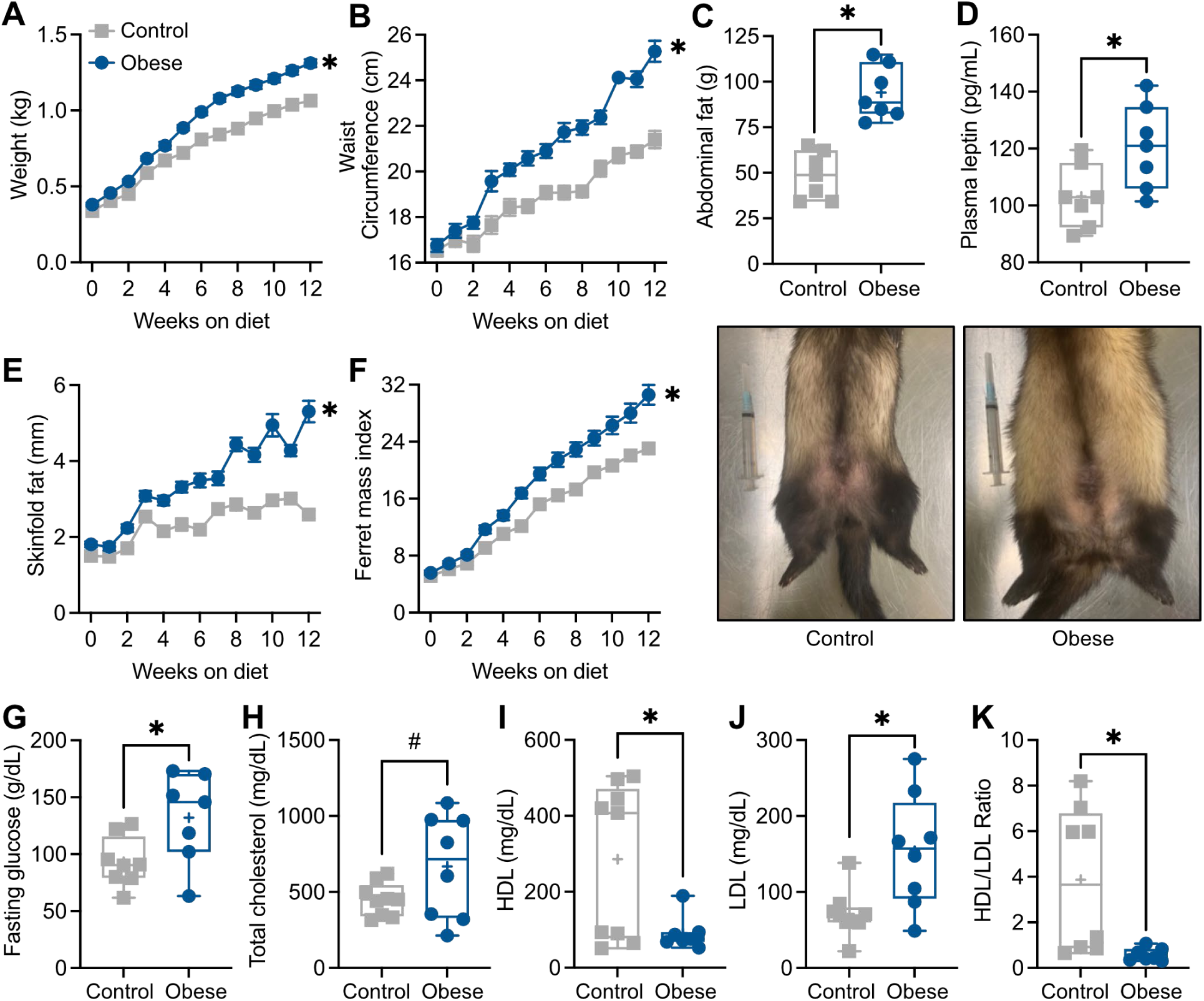
Inducing obesity and metabolic syndrome in ferrets. Male ferrets (6 weeks of age) were placed on control or high calorie diet and monitored for 12 weeks. (A) Weight (*p*<0.0001), (B) waist circumference (*p*<0.0001), and (C) skinfold fat (*p*<0.0001) were assessed weekly. (D) Ferret mass index was calculated from physical measurements (*p*<0.0001). Data was analyzed by 2-way ANOVA with repeated measures (*p*-value represents simple main effect of diet). (E) Animals were sacrificed post-diet and abdominal fat deposits removed and weighed (*p*<0.0001 by unpaired *t* test). (F) Plasma leptin levels post-diet were determined by ELISA (*p*=0.0278 by unpaired *t* test). (G) Differences in physical appearance of control and obese ferrets. (H) Fasting glucose (*p*=0.0309), (I) total cholesterol (*p*=0.0878), (J) high-density lipoprotein (HDL) (*p*=0.0265), and (K) low-density lipoprotein (LDL) (*p*= 0.0079) levels in the plasma of ferrets post-diet. (L) Ratio of HDL to LDL (*p*=0.0187). Significance of (H-L) was determined by unpaired *t* test. Data represents 2 independent experiments of n=7-10/group. Error bars represent standard error of the mean (SEM) (A-D) or minimum value to maximum value (E-F, H-L). \**p*<0.05, ^#^*p*<0.10.

**Fig. S2.**
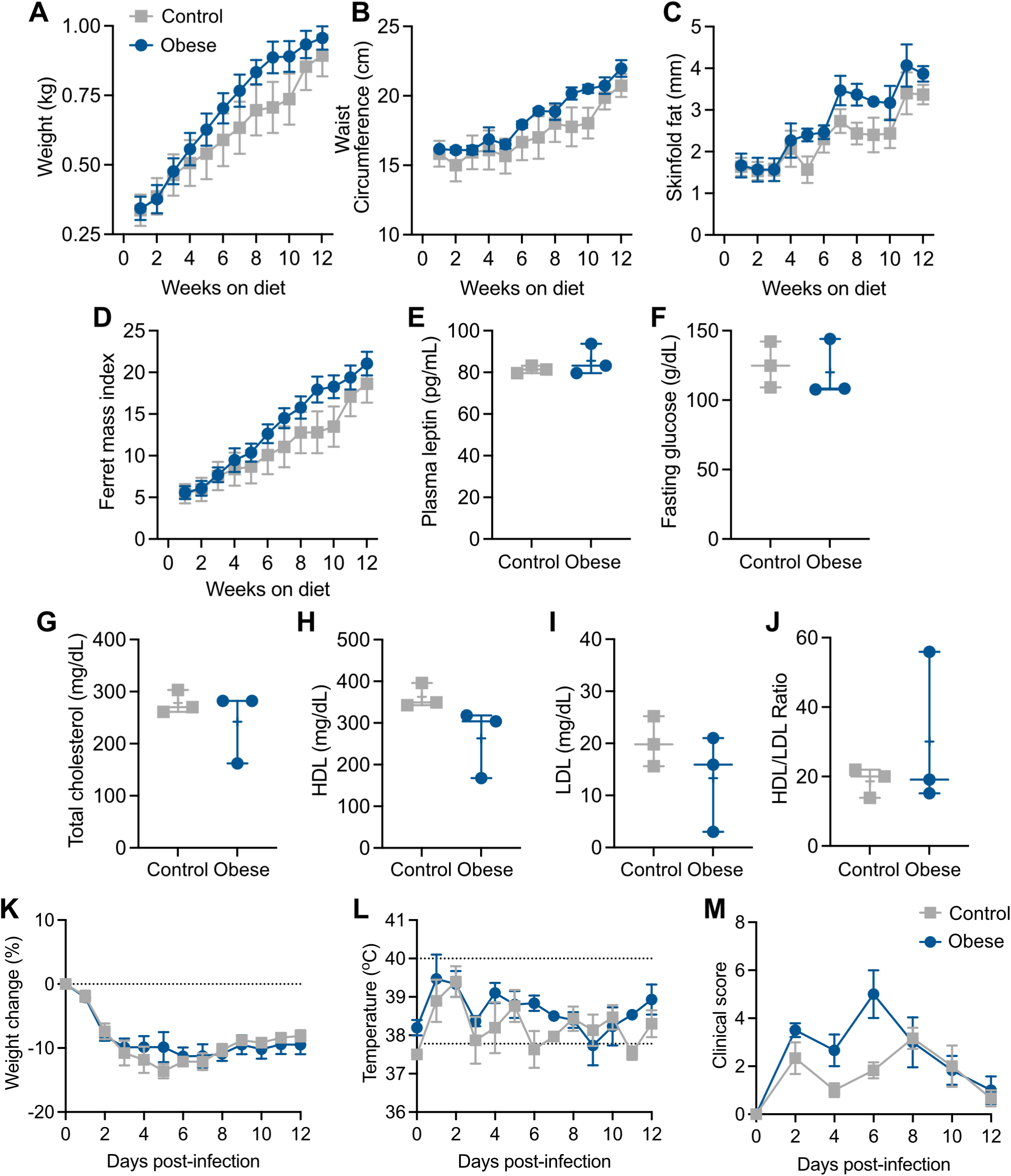
Obesity and disease severity in female ferrets. Female ferrets (6 weeks of age) were placed on control or obese diet and monitored for 12 weeks. Weekly measurements were taken of (A) weight (*p*=0.4308), (B) circumference (*p*=0.3888), and (C) skinfold fat (*p*=0.2389). (D) Ferret mass index was calculated from physical measurements (*p*=0.3995). Data was analyzed by 2-way ANOVA with repeated measures (*p*-value represents simple main effect of diet). (E) Leptin (*p*=0.3987), (F) fasting glucose (*p*=0.7424), (G) total cholesterol (*p*=0.4396), (H) high-density lipoprotein (HDL) (*p*=0.1222), and (I) low-density lipoprotein (LDL) (*p*=0.3166) levels in the plasma of female ferrets post-diet. (J) Ratio of HDL to LDL (*p*=0.4401). Data shown in E-J was analyzed by unpaired *t* test. (K) After 12 weeks on diet, female ferrets were infected with 10^6^ TCID_50_ A/California/04/2009 (H1N1) virus and monitored for 12 days. Weight (*p*=0.9096), (L) body temperature (*p*=0.3400), and (M) clinical scores (*p*=0.1703), were recorded throughout infection. Significance was determined by 2-way ANOVA with repeated measures (*p*-value represents simple main effect of diet). Data represents 1 independent experiment of n=3/group. Error bars are standard deviation of the mean (SD) and dashed lines indicate baseline weight (K), normal range (L) or lower limit of detection (M). \**p*<0.05, ^#^*p*<0.10.

**Table S1.**
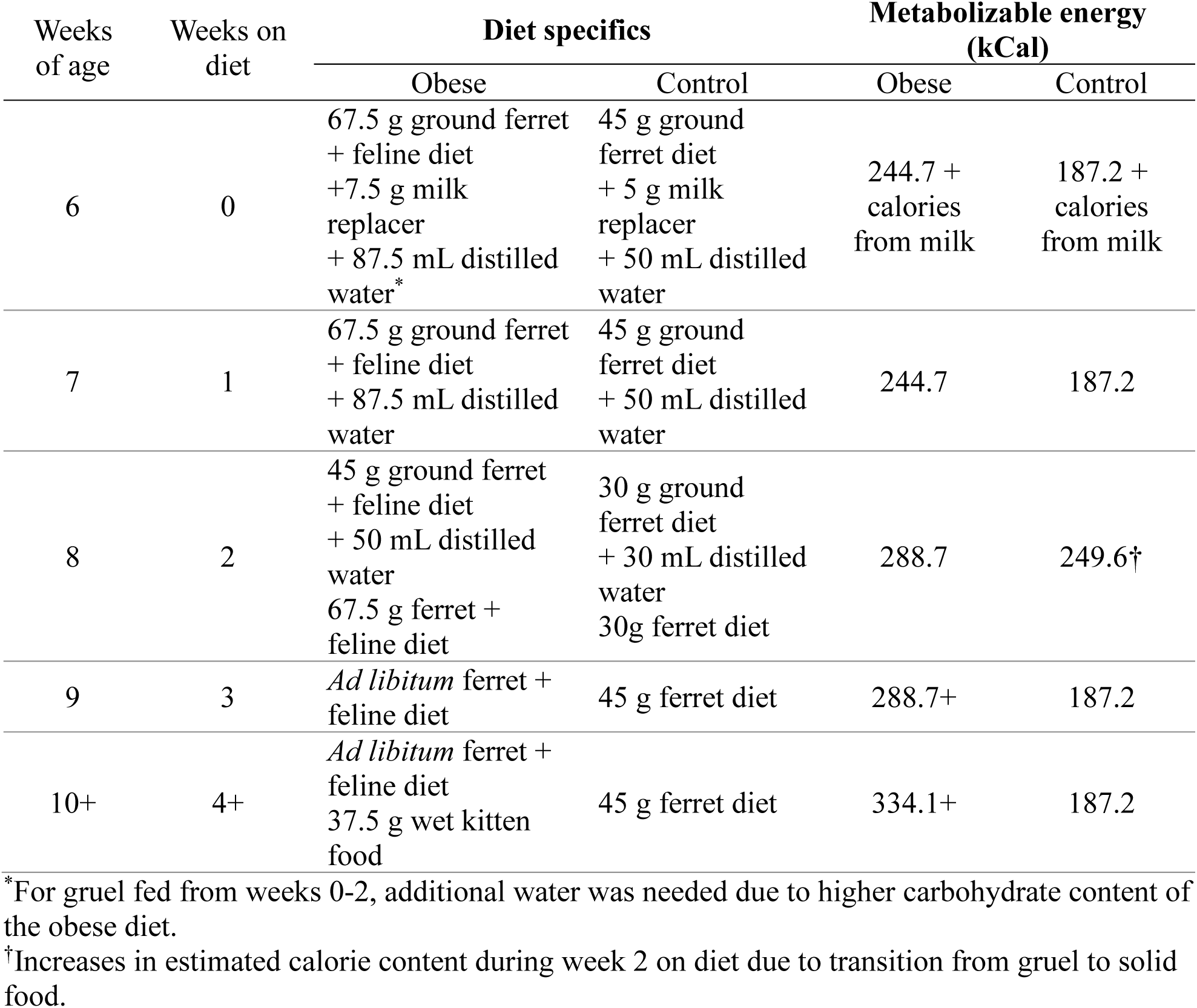
Diet-induced obesity feeding plan per ferret.

**Table S2.**
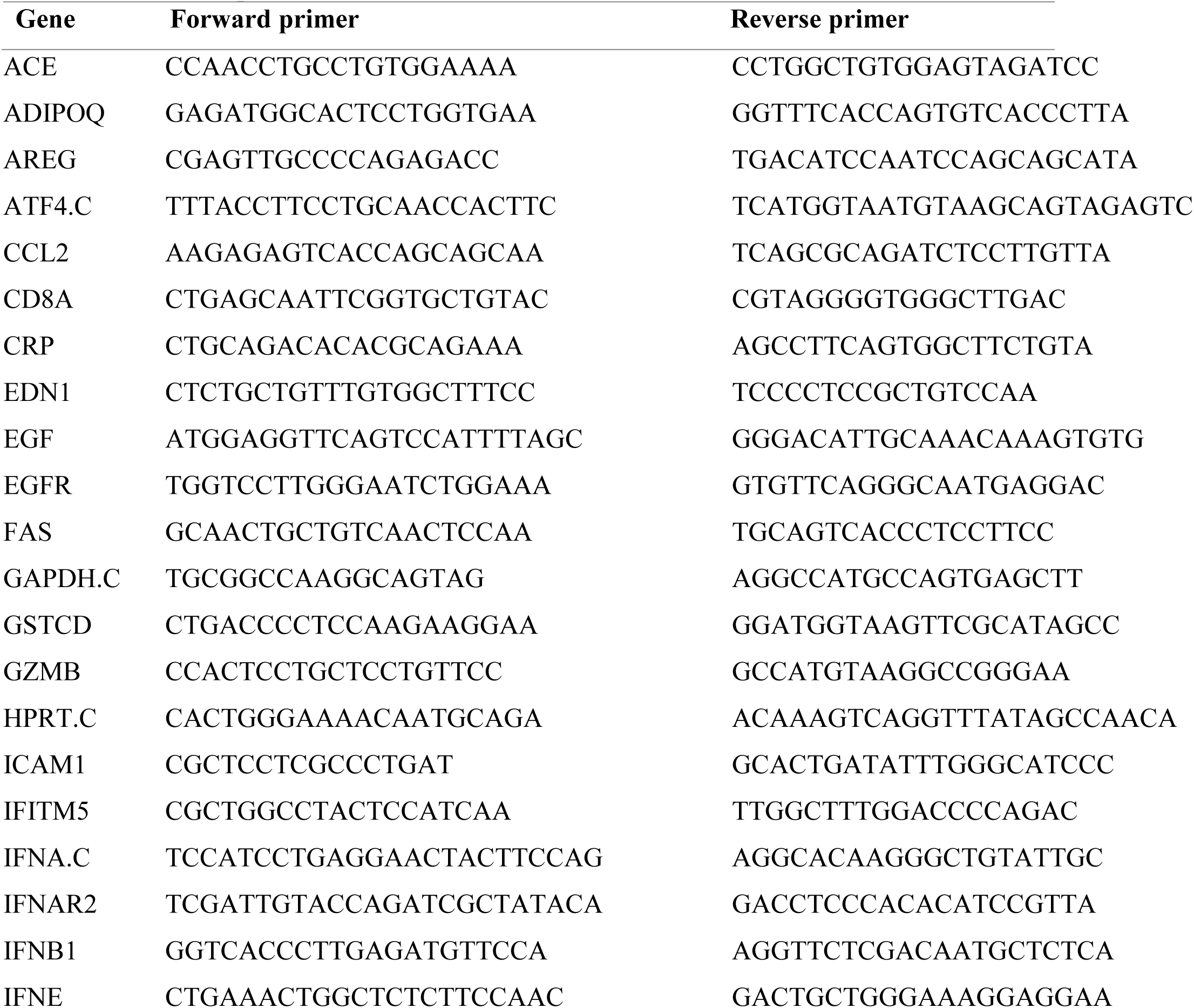
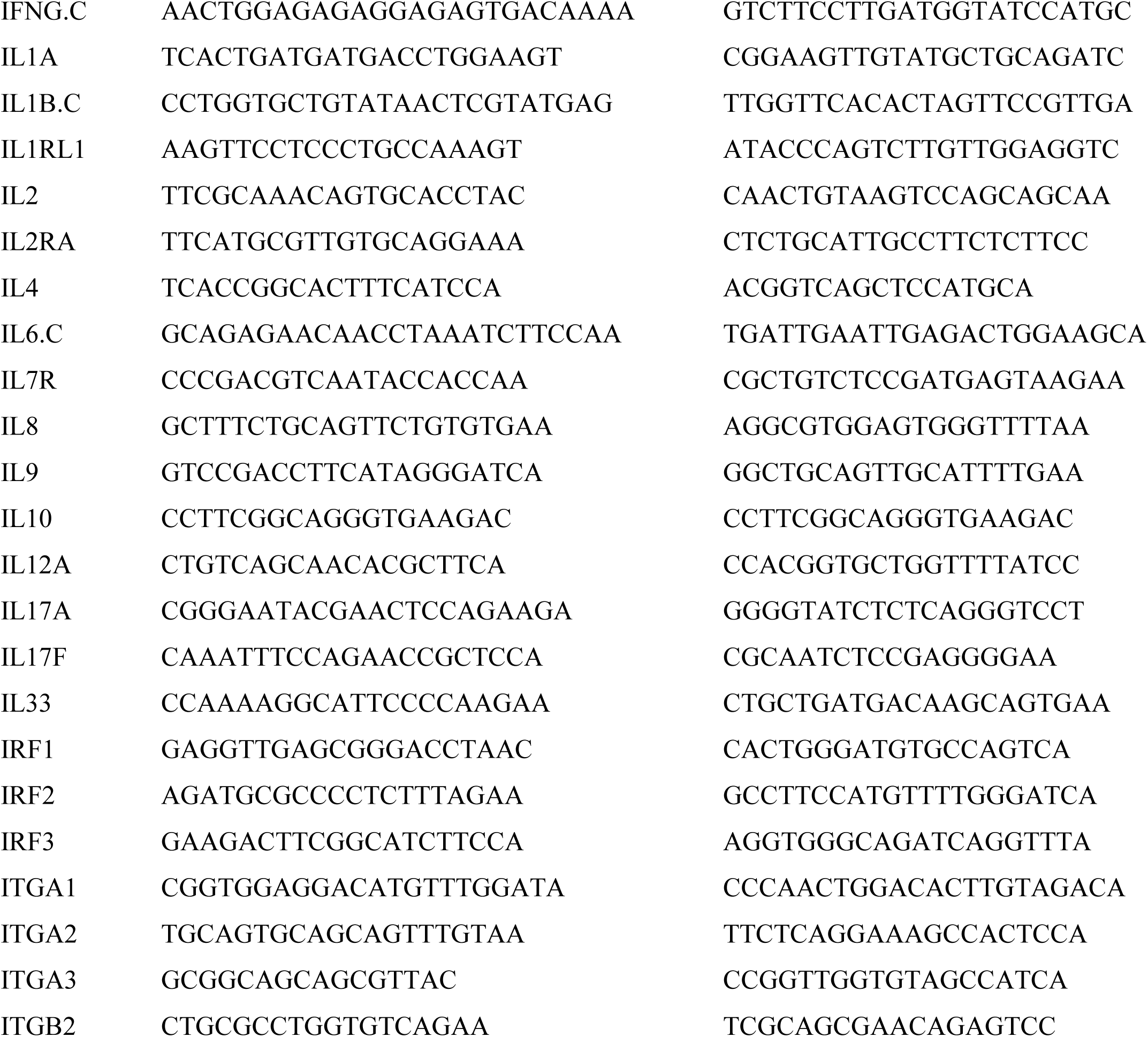
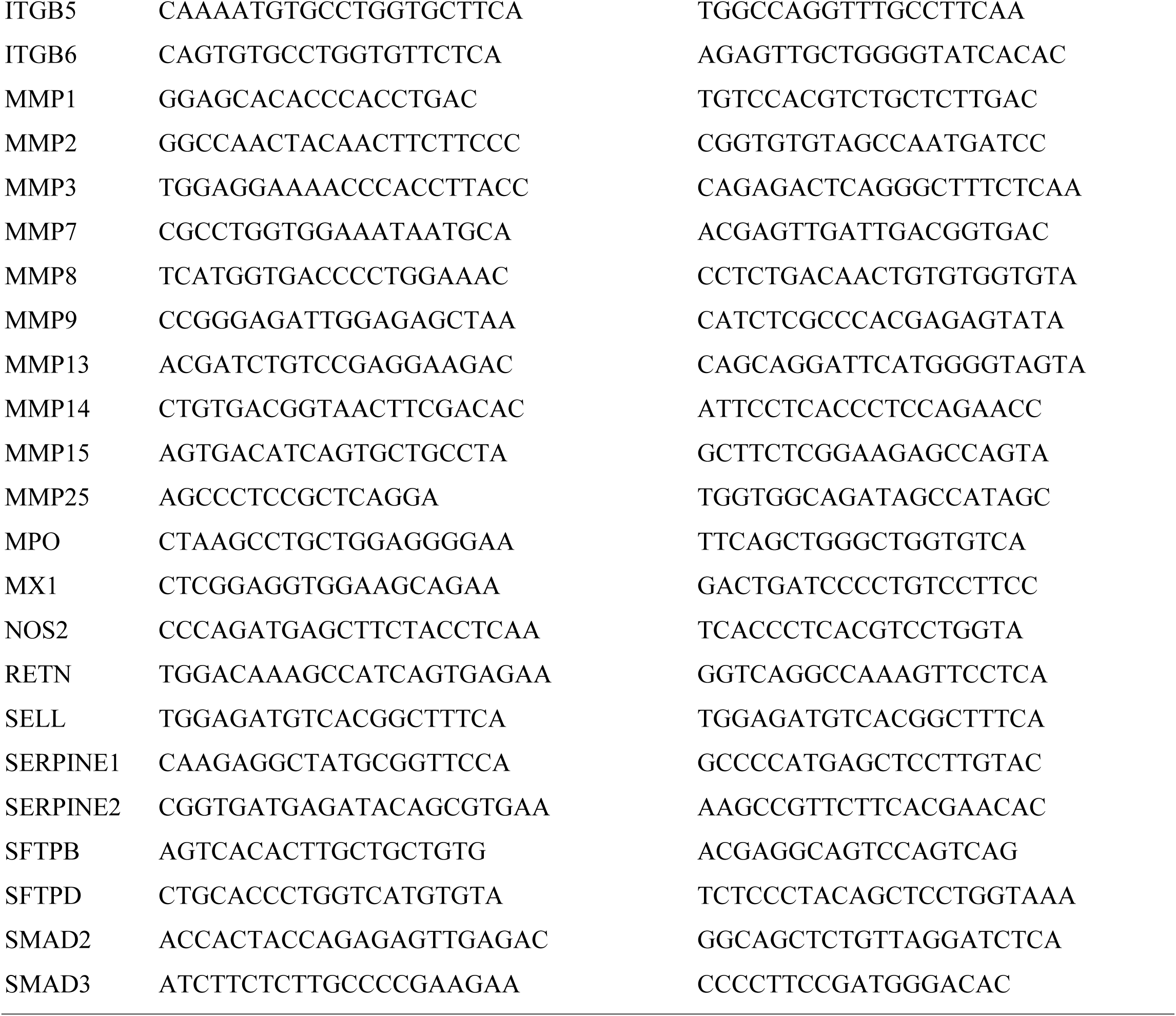
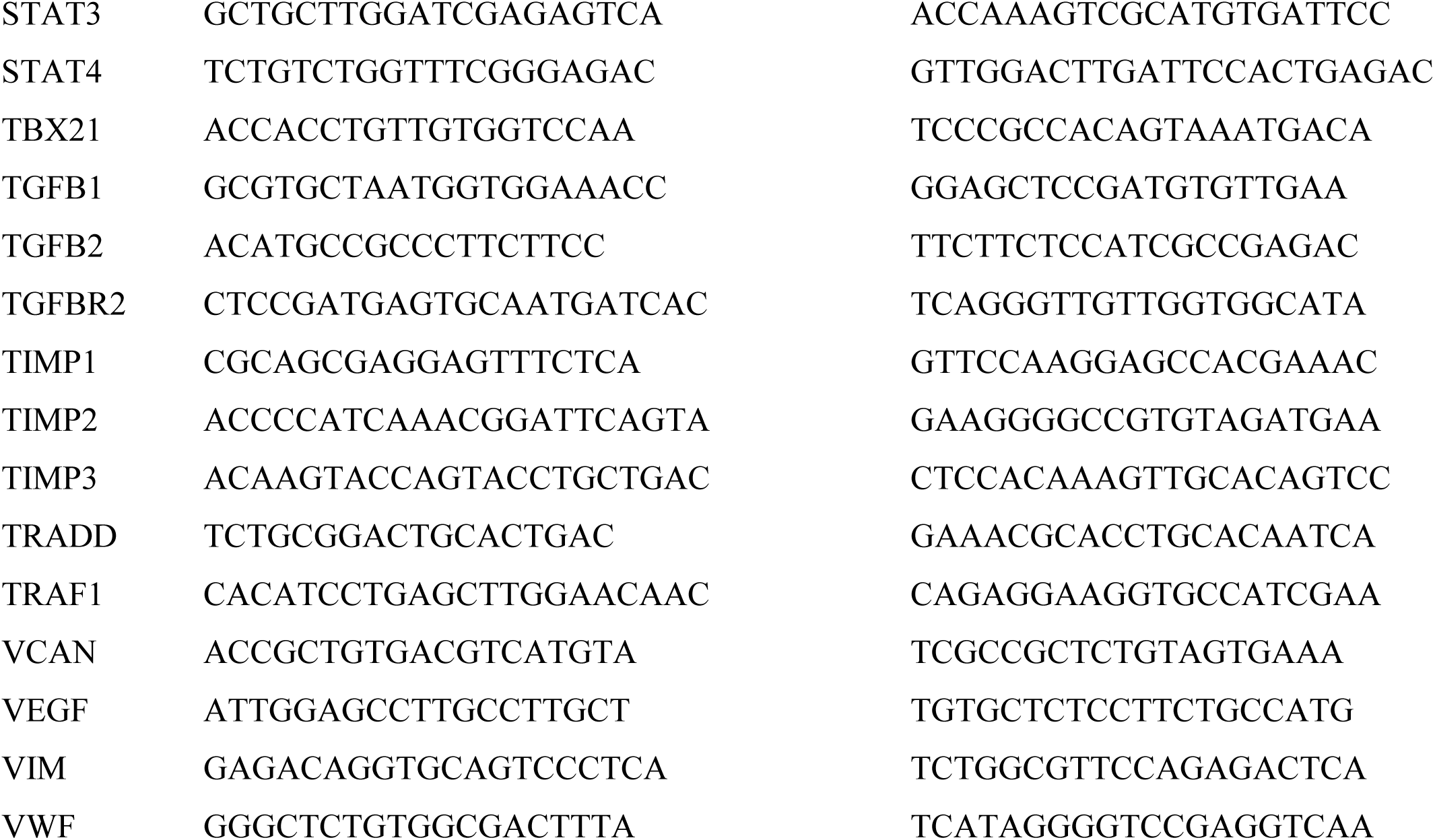
Primer sequences.

